# Improving integrative 3D modeling into low- to medium- resolution EM structures with evolutionary couplings

**DOI:** 10.1101/2021.01.14.426447

**Authors:** Caitlyn L. McCafferty, David W. Taylor, Edward M. Marcotte

## Abstract

Electron microscopy (EM) continues to provide near-atomic resolution structures for well-behaved proteins and protein complexes. Unfortunately, structures of some complexes are limited to low- to medium-resolution due to biochemical or conformational heterogeneity. Thus, the application of unbiased systematic methods for fitting individual structures into EM maps is important. A method that employs co-evolutionary information obtained solely from sequence data could prove invaluable for quick, confident localization of subunits within these structures. Here, we incorporate the co-evolution of intermolecular amino acids as a new type of distance restraint in the Integrative Modeling Platform (IMP) in order to build three-dimensional models of atomic structures into EM maps ranging from 10-14 Å in resolution. We validate this method using four complexes of known structure, where we highlight the conservation of intermolecular couplings despite dynamic conformational changes using the BAM complex. Finally, we use this method to assemble the subunits of the bacterial holo-translocon into a model that agrees with previous biochemical data. The use of evolutionary couplings in integrative modeling improves systematic, unbiased fitting of atomic models into medium- to low-resolution EM maps, providing additional information to integrative models lacking in spatial data.

## Introduction

The era of high throughput -omics data has transformed our understanding of systems biology through the interrogation of big data. The generation of this data has led to an abundance of information in databases. UniProt, for example, is flooded with over 180 million sequences, most of which still lack complete structures or annotations^1^. Furthermore, advancements in algorithmic approaches have illuminated the complexity of information within a single protein sequence^2-4^, and how, when aligned with 100s of other sequences, can produce a prediction of functional sites or even a family of proteins^5-7^. The use of this co-evolutionary information in a protein sequence allows us to scour the depths of these databases to uncover novel patterns that lead to robust biological hypotheses. An exciting recent example of such an approach is direct coupling analysis applied to residue co-evolution^8^.

Residue co-evolution as it applies to structural biology is based on the idea that amino acids close in three-dimensional (3D) space within a protein structure exhibit statistical coupling through the course of evolution, such that mutations in one may occasionally be accompanied by mutations to the other to compensate for the change without altering the 3D structure^9^. These couplings between nearby residues can be distinguished based on analysis of primary amino acid sequences in multiple sequence alignments and used to predict 3D structures from sequence information alone^10; 11^. Not only are evolutionary couplings being used as a standalone structural predictor, but they are also being used in various machine learning algorithms to predict structure from sequence. For example, Alphafold won the CASP13 and CASP14 competition where they incorporated evolutionary data into their deep learning approach of *ab initio* protein structure prediction^12^. Moreover, when combined with other pairwise distance restraints, such as sparse NMR data from larger proteins, evolutionary couplings can be used to build more accurate and complete protein structures^13; 14^. The dependency between residues that evolutionary coupling relies on has also been used to predict mutation effects^15^. This differs from conservation in that it considers epistasis by modeling interactions of all residue pairs, allowing for the quantification of multiple mutations. As a result, this method can better capture experimental mutational fitness landscapes.

Beyond intramolecular patterns, evolutionary coupling has been used to detect intermolecular interactions. This has been demonstrated between amino acids and nucleotides as well as nucleotide/nucleotide interactions in RNA-protein complexes^16^. It has also been used to predict protein-protein interactions^17; 18^. Moreover, combined methods of disentangling evolutionary couplings have been applied across proteomes^19; 20^.

When combined with other experimental techniques, evolutionary couplings have the ability to improve structural predictions. The Integrative Modeling Platform^21^ can use distance restraints, including chemical crosslinks, along with atomic models and electron microscopy data to produce predictions of large molecular ensembles^22; 23^. These methods use curated scoring functions combined with Monte Carlo sampling and simulated annealing to produce ensembles of models that satisfy experimental restraints. The ensemble is then evaluated *via* statistical testing and validated against biological data such as genetic or biochemical experiments. Additional distance restraints derived from evolutionary coupling data have the potential to provide useful information for systems with few crosslinks and could potentially be used as the only distance restraints when no other data are available.

During interpretation of low-resolution EM maps, the reconstruction is often treated as an envelope where crystal structures or homology models can be docked into regions following either segmentation^24^ or localization of specific tags within the map^25^. While these methods are effective in identifying an area of the low-resolution map where a protein may be located, they do not provide higher resolution information such as residue-residue interactions that may give insight into orientation and interaction interfaces on the individual protein subunits. The combination of higher resolution data such as information from amino acid evolutionary couplings with lower resolution EM maps in integrative modeling could produce an objective and unbiased configuration of protein subunits that satisfy residue-residue distance restraints derived from evolutionary couplings and protein complex shape from EM. Here, we focus on developing a distance restraint for evolutionary couplings to be used when fitting atomic structures into low-resolution EM maps.

We demonstrate that information from evolutionary couplings can be used as a distance restraint in the integrative modeling of atomic structures into low-resolution electron microscopy maps. We present a method for selecting intermolecular couplings to be used for integrative modeling based on internal structural controls as well as a weighting of couplings that is tolerant of false-positive pairs. We show that a lack of distance restraints from evolutionary couplings results in a poor structural prediction with the quinol-fumarate reductase complex. We investigate evolutionary couplings in dynamic protein complexes that experience conformational changes within the macromolecular machine and within individual subunits using the BAM complex. Finally, we build an integrative model of the bacterial holo-translocon using evolutionary couplings between the 6 subunits, a 14 Å cryo-EM map, and a combination of K-ray crystal structures and high-confidence homology models. We validate our model against another independently derived model as well as extensive data regarding the mechanism, chemistry, and physics of the complex.

## Materials and methods

### Protein Complex Selection

We used the browse stoichiometry search in the Protein Data Bank to select *E. coli* complexes with stoichiometries ABC, ABCD, and ABCDE. Complexes for modeling were selected based on Neff rating and shape of the coupling plot. For example, complexes without high scoring intramolecular couples cannot identify intermolecular couples. Information on the complexes modeled can be found in **Supplemental Table 1**.

### Evolutionary Coupling Analysis

We computed the evolutionary coupling analysis using the EV coupling python framework^26^. The sequence search was performed using the uniref100^27^ database. We used the best_hit protocol for the sequence alignment, which pairs the sequences that have the highest percent identity to the target sequence for each genome. The configuration file with all of the run parameters is available on the github page: https://github.com/marcottelab/CoEVxIMP. We do an all-by-all coupling analysis for all subunits in the protein complex. For example, if the complex has subunits ABC, we calculate couplings between 3 groups: AB, AC, and BC. Likewise, ABCDE couplings are calculated between 11 groups. A complex of size N has couplings between N! groups. For couplings between each protein pair, we take the top 50,000 coupling scores (cn scores) of all residue pairings; beyond this, the couplings would not pass any selected threshold. We select intermolecular couplings based on a cutoff cn score, which is calculated based on a 10-coupling, moving average of intramolecular distances. We took the cn score when the moving average first exceeded the following three distances: 10 Å, 15 Å, and 20 Å. We calculated the distances between Cβ atoms because previous studies have shown that side-chain atoms are more likely to be structurally coupled than backbone atoms^28^. Cβ atoms that were not present in the structure such as the case with missing amino acids or glycine residues were removed from the moving average calculation.

### Evolutionary Coupling Analysis of Stoichiometric Complexes

We handle more complex stoichiometries, such as A2B in PDB: 4I98, by separating intramolecular contacts from intermolecular contacts of the same protein. We calculate a rolling mean and standard deviation of intramolecular distance across 100 coupling increments. We use the moving average of 100 couplings so that there is a larger sample size for the standard deviation calculation. Any distance that was greater than *x̄* ± 2*s* and cn score greater than the 10-Å cutoff, c_0_, was determined to be an intermolecular coupling between identical subunits. These couplings were then removed from the intramolecular internal calibration and a new 10-Å cutoff, c_1_, was calculated. The process was then repeated to identify more intermolecular contacts using the new cutoff until c_n_ remains constant.

### IMP Integration

We selected complexes for integrative modeling with IMP (Integrative Modeling Platform) based on the signal from evolutionary couplings. Complexes that showed no correlation within intramolecular contacts i.e. low-scoring intramolecular pairs that are close in distance are not reliable for intermolecular interpretation. The low signal can be a result of not having enough sequences or even simply too much variation in sequences. We selected 5 protein complexes that fit these criteria, PDB IDs: 1FFT, 1L0V, 5MRW, 5D0O, and 5D0Q.

We used the Integrative Modeling Platform python modeling interface^29^ to assemble the protein complexes using atomic structures, excluded volume, sequence connectivity, a synthetic EM map, and the evolutionary coupling distance restraints (**Figure 2**). The atomic structures were taken from the PDB of the protein complex and treated as rigid bodies, while the simulated EM map was created by low pass filtering the PDB to 10 Å using gmcovert^30^. The maps were then approximated using a Gaussian mixture model of 50 components^30^. Groups of 20 residues of each subunit were also approximated by Gaussian components. The evolutionary couplings restraints were generated using the method described above.

Each protein complex was subjected to three different modeling runs. The first used intermolecular interactions predicted from the 10-Å score cutoff couplings, the second used 15-Å score cutoff couplings, and the third used the 20-Å score cutoff couplings. The minimum distance for the restraint was set to 0 Å and the maximum distance was the cutoff score distance used plus one standard deviation. Each residue pairing distance restraint was weighted as 10 x cn score. All runs were optimized using Monte Carlo sampling and score based on excluded volume, sequence connectivity, evolutionary coupling distance, and EM map. **Table 1** describes run parameters used for each of the complexes.

**Table 1.** Parameters used to model complexes.

**Figure 6: Integrative modeling of the bacterial holo-translocon**
| PDB | Cutoff threshold (Å) | Number of couplings | Number of runs | Frames per run | Cluster size | Cluster precision (Å) |
| --- | --- | --- | --- | --- | --- | --- |
| 1FFT | 10 | 19 | 4 | 10,000 | 31,349 | 11.255 |
| 1FFT | 15 | 39 | 6 | 10,000 | 32,767 | 9.369 |
| 1FFT | 20 | 63 | 6 | 10,000 | 32,393 | 11.526 |
| 1L0V | 10 | 14 | 10 | 10,000 | 1,600 | 30.000 |
| 1L0V | 15 | 54 | 10 | 10,000 | 9,884 | 15.000 |
| 1L0V | 20 | 93 | 10 | 10,000 | 5,585 | 19.000 |
| 5D0O | 10 | 45 | 10 | 10,000 | 2,053 | 9.000 |
| 5D0O | 15 | 165 | 10 | 10,000 | 1,973 | 28.000 |
| 5D0O | 20 | 202 | 10 | 10,000 | 6,500 | 27.000 |
| 5D0Q | 10 | 31 | 10 | 10,000 | 25,007 | 21.672 |
| 5D0Q | 15 | 47 | 10 | 10,000 |  |  |
| 5D0Q | 20 | 123 | 10 | 10,000 | 38,753 | 17.632 |
| 5MRW | 10 | 90 | 5 | 10,000 | 31,258 | 1.100 |
| 5MRW | 15 | 140 | 10 | 10,000 | 13,271 | 2.151 |
| 5MRW | 20 | 207 | 10 | 10,000 | 29,850 | 1.099 |

### Validation

We validated our models using the sampling exhaustiveness approach^31^. Good scoring models were selected using hdbscan clustering^32^. We then selected the cluster that had both a high score based on satisfying input restraints and a large number of models. Assessing sampling exhaustiveness of our model involved testing the convergence of our high-scoring models from the selected cluster, confirming the similarity of the score distribution between the two samples, testing that the models in the structural clusters were proportional to each cluster’s size, and finally identifying structural similarity between models from each sample. These tests are quantified using sampling precision, Cramer’s V test, and computing localization densities for each subunit. To validate our models against the input data, we used a cross-correlation coefficient between the EM map used in the modeling and the probability density to further evaluate the predicted models and select which evolutionary coupling threshold performed the best.

### Modeling the bacterial holo translocon

The bacterial holo translocon (HTL) is made up of 7 subunits: SecY, SecE, SecG, YidC, SecD, SecF, and YajC. Our modeling includes all subunits with the exception of YajC due to the small size of YajC and its suspected instability within the HTL complex. Full-length YidC (P25714), SecD (P0AG90), SecE (P0AG96), and SecG (P0AG99) were modeled using I-TASSER^33^. The *E. coli* YidC crystal structure (PDB ID: 6AL2)^34^ and the homology model of Sec F from PDB ID: 5MG3^35^ were used. SecY and SecF were treated as rigid bodies; SecE, SecD, and YidC were treated as chains of rigid bodies; and SecG was treated as a combination of rigid bodies held together with beads, to allow flexibility within the peptide.

We used the 10-Å score cutoff for the selection of intermolecular couplings. In total 15 couplings were used between 6 subunit pairs, in which each subunit had at least one coupling to another subunit. Each coupling was weighted according to the method described above. We used the 14-Å resolution cryo-EM map (EMDB-3506) for our modeling of the HTL. The map was approximated by a Gaussian mixture model (GMM) of 50 components. Each atomic model was approximated by a GMM for every 20 residues. 200,000 models were computed from 10 initial positions. Clustering based on scores gave a high-scoring cluster of 10,000 models that satisfied the restraints used in the sampling. Clustering the models selected gave a sampling precision of 37 Å with 90.27% of the high-scoring models included in the final cluster.

## Results

We were interested in utilizing co-evolutionary information that is available in the form of sequence data to guide the modeling of molecular assemblies. Specifically, we wished to develop a systematic method for fitting atomic structures into medium- to low-resolution electron microscopy maps using information from evolutionary coupling data to illuminate subunit orientation and protein-protein interaction sites. To do this, we focused on building a distance-based restraint for evolutionary couplings into the Integrative Modeling Platform (IMP) that can use several lower-scoring couplings but is still tolerant of some false-positive pairs.

### Calibrating intermolecular couplings based on internally-consistent intramolecular couplings

The determination of a score cutoff is essential when making intermolecular contact predictions based on evolutionary coupling data. This is due, in part, to the nature of the evolutionary coupling analyses that look at every pair of amino acids, both intra- and intermolecular, and assign a score based on the assumption that most pairs are not coupled, and those that are, are outliers that lie in the tail of the distribution^17^. Previous studies have suggested different score cutoffs^17; 36; 37^, however, we were interested in leveraging the information contained in a known individual atomic structure of a protein to guide the intermolecular coupling score cutoff. For this reason, we use known X-ray crystal structures (or a high confidence homology model) as an internal control for selecting intermolecular contacts. This is done by plotting the cn score against distance for the intramolecular contacts for each protein pair being analyzed. We then compute a moving average for groups of 10 couples and choose the score cutoff as the point that the moving average reaches 10, 15, and 20 Å (described in **Materials and methods**), ultimately testing all-by-all residue couplings across all subunit pairs. A benefit of this method is that it recalibrates the score cutoff for each protein pair, allowing flexibility with the score cutoff, as it reflects the data used and, in some cases, extracting lower-scoring data.

We demonstrate the utility of this internal calibration approach with the *E. coli* ubiquinol oxidase complex (**Figure 1**). The intramolecular contacts that are mapped onto the crystal structures (**Figure 1A**) are used to set the score cutoff for the intermolecular contacts mapped onto the known complex structure (**Figure 1B**). It is evident from the known structure that subunits B and C do not contact each other, and this is supported by the graph of the cn score against the distance (**Figure 1C**). The intermolecular couplings that are selected using the internal calibration help to stitch the subunits together to form the complex.

**Figure 1:**
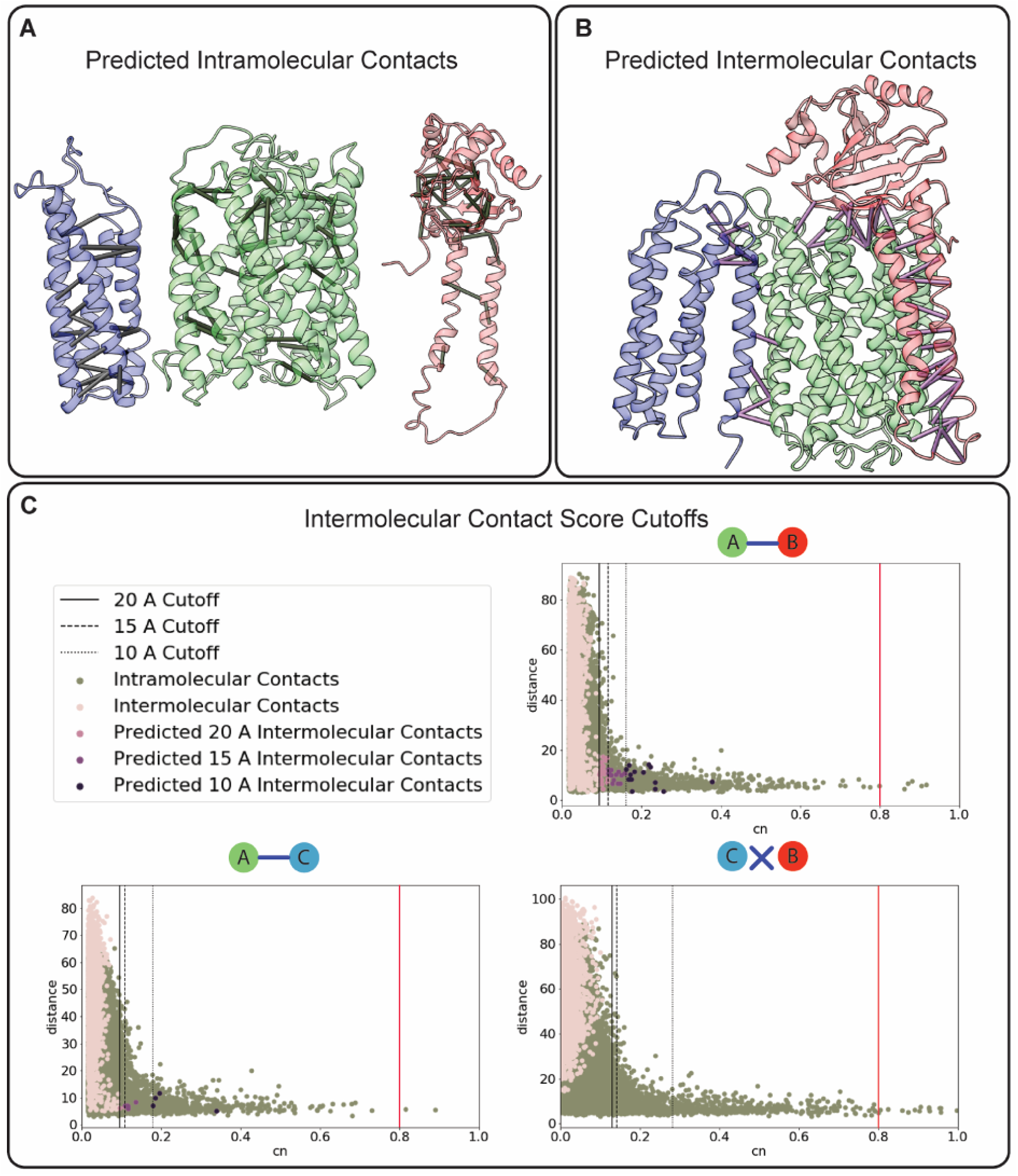
Selecting intermolecular evolutionary couplings. Evolutionary couplings are computed between amino acid pairs using the EVcouplings python package^26^. Intermolecular couplings are selected using a score cutoff determined by an internal control, the intramolecular coupling distances of each individual subunit in the complex. **A**. Intramolecular couplings selected based on a score cutoff determined from the moving average of Cb distances (< 10 Å) between amino acids shown on the subunits of ubiquinol oxidase (PDB: 1FFT). **B**. The score cutoff determined from the internal control in **A** was used to select intermolecular couplings shown on the ubiquinol oxidase complex structure (PDB:1FFT). **C**. Intramolecular moving average distances of 10, 15, and 20 Å were used to determine different score cutoffs for intermolecular couplings for each pairing of the subunits. The red line shows the recommended score for selecting intermolecular couplings^17^.

### Inter- vs Intra-molecular couplings can be distinguished within homomeric assemblies

As protein assemblies often involve complex stoichiometries, we specifically tested if a variant of this general approach could distinguish intermolecular coupling between homomeric interaction partners from intramolecular couplings within the proteins. To do this we considered all intramolecular couplings that are calculated to fall outside of the distance of two standard deviations above the moving average and score greater than the calculated 10-Å cutoff score. This is repeated until there is no change in the cutoff score (described in **Materials and methods**). We hypothesized that this method could be used to identify both the obvious outliers and more subtle contacts.

We tested this approach by applying it to the symmetric condensin Smc homodimer in the context of their interaction with the protein kleisin, forming a 2:1 Smc:kleisin assembly^38^ (PDB: 4I98) (**Figure 2**). We plotted distances for intramolecular contacts that were selected using this method and the changing 10-Å cutoff score (**Figure 2A**). When those intramolecular distances for the selected outlier are computed between the identical subunits rather than within a subunit, the distance decreases for 92% of the couplings selected (**Figure 2B**). Thus, the internal calibration of intramolecular couplings on a 3D structure can help determine if an intramolecular contact truly lies within the protein or instead comprises an intermolecular contact between copies of the protein.

**Figure 2:**
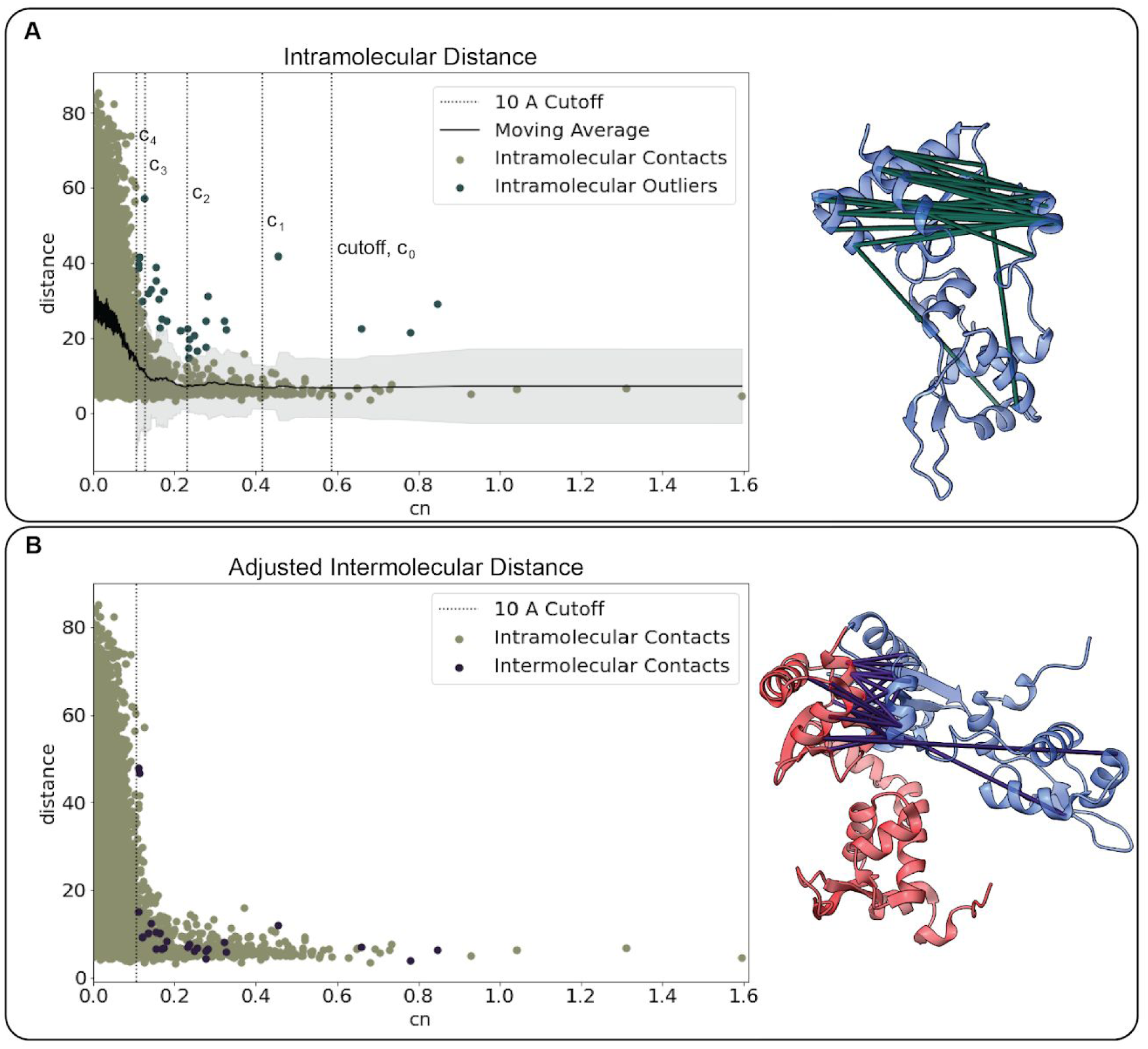
Disentangling intramolecular couplings from intermolecular couplings of homomeric interaction partners. Intramolecular couplings were computed and plotted against their distance in the reference protein X-ray crystal structure (PDB: 4I98). **A**. The moving average is shown with 2 standard deviation distance above and below (light grey). The intramolecular couplings with a cn score greater than the 10-Å cutoff score with distances above the grey region are shown in dark green. After removal of the outliers, the 10-Å cutoff score is recomputed (c_0_, c_1_, c_2_, c_3_, c_4_, c_5_). The intramolecular couplings removed are illustrated on the x-ray crystal structure. **B**. The distance of the selected intramolecular couplings from **A** are recomputed as intermolecular distances between the identical subunits. These coupling are again illustrated on the x-ray crystal structures showing the smaller distance between the subunits.

### An evolutionary coupling restraint for integrative modeling

Incorporating the evolutionary coupling data into IMP involves expressing the pairs as a weighted distance restraint. We use our internal control method to select the score cutoff for each protein pair. Each of the couplings is represented as a basic distance restraint in IMP, where it is weighted based on the cn score, weighting high scoring couples more heavily in the model scoring. It is worth mentioning that while the lower score cutoffs may increase false-positive contacts, this is balanced by the weighting of each coupling, and the distribution outliers will still be given a larger weight in the scoring function. To evaluate this method, we tested the couplings determined for three different bacterial complexes (PDBs: 1FFT, 1L0V, 5MRW) and one dynamic complex (PDBs: 5D0Q, 5D0O) using the IMP scheme (**Figure 3**). To create the simulated EM map, each of the complexes was low-pass filtered to a resolution of 10 Å. Ubiquinol oxidase (PDB: 1FFT) and KdpFABC (PDB: 5MRW) were heavily coupled (**Supplementary Data**) and as a result produced robust predictions with clustering precisions of <11.5 Å and <2.2 Å, respectively (**Figure 4A**).

**Figure 3:**
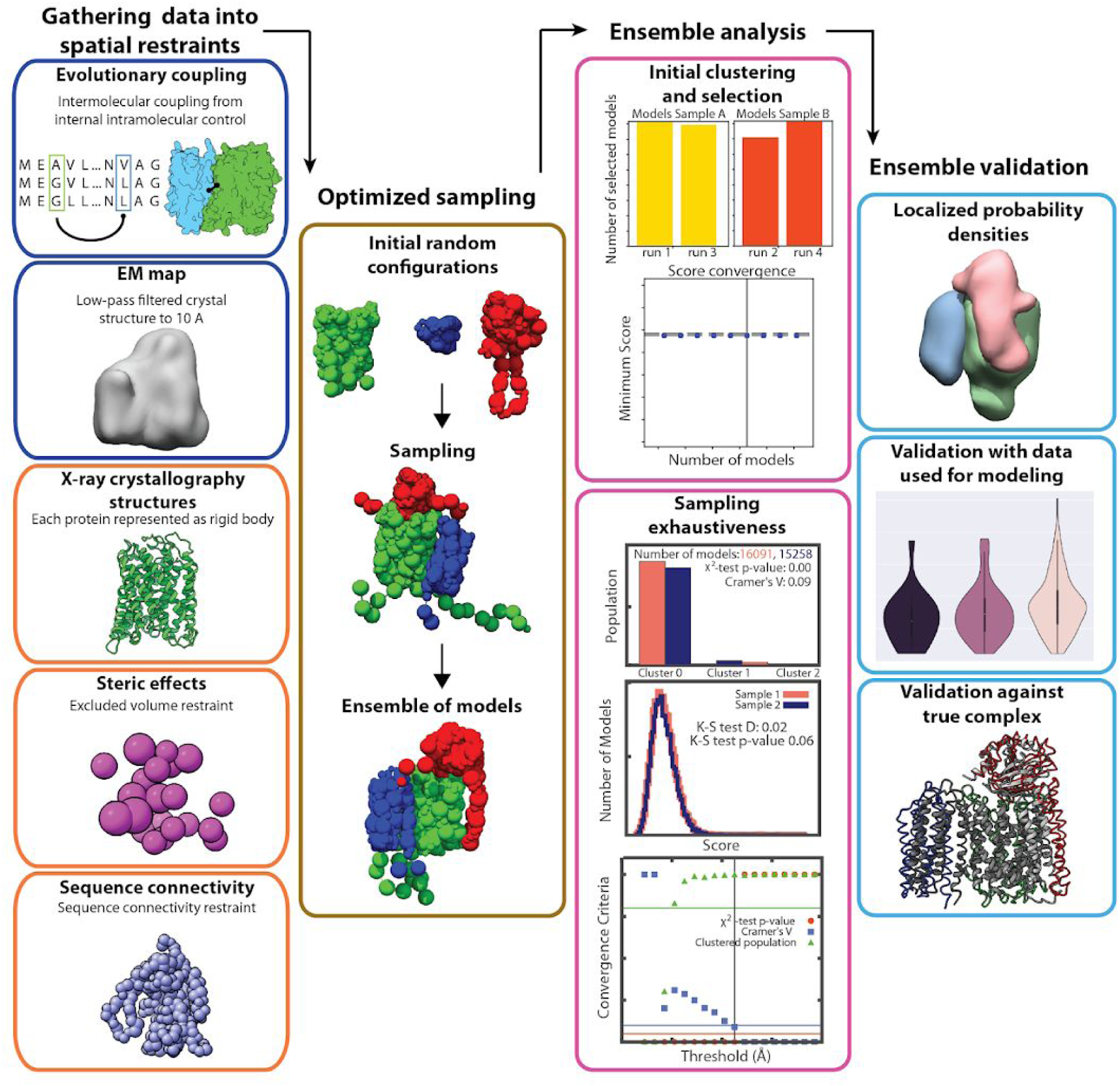
Integrative Modeling Platform (IMP) scheme for validating the evolutionary coupling restraint. Following the IMP protocol^20^, data is converted into spatial restraints. The evolutionary coupling restraint is included as a distance restraint and weighted by its coupling score. The dark blue boxes represent experimental data, in this case evolutionary couplings and a synthetic 10-Å resolution EM map. The orange boxes show crystal structures, statistical inferences, and physical properties. The gold box displays the sampling and scoring of the models. The models are analyzed via an initial clustering to select a high scoring group of models from various simulation runs followed by sampling exhaustiveness^29^ (pink boxes). The light blue boxes show the ensemble validation against the data used in the modeling, the probability density for each subunit, and a comparison against the known structure.

**Figure 4:**
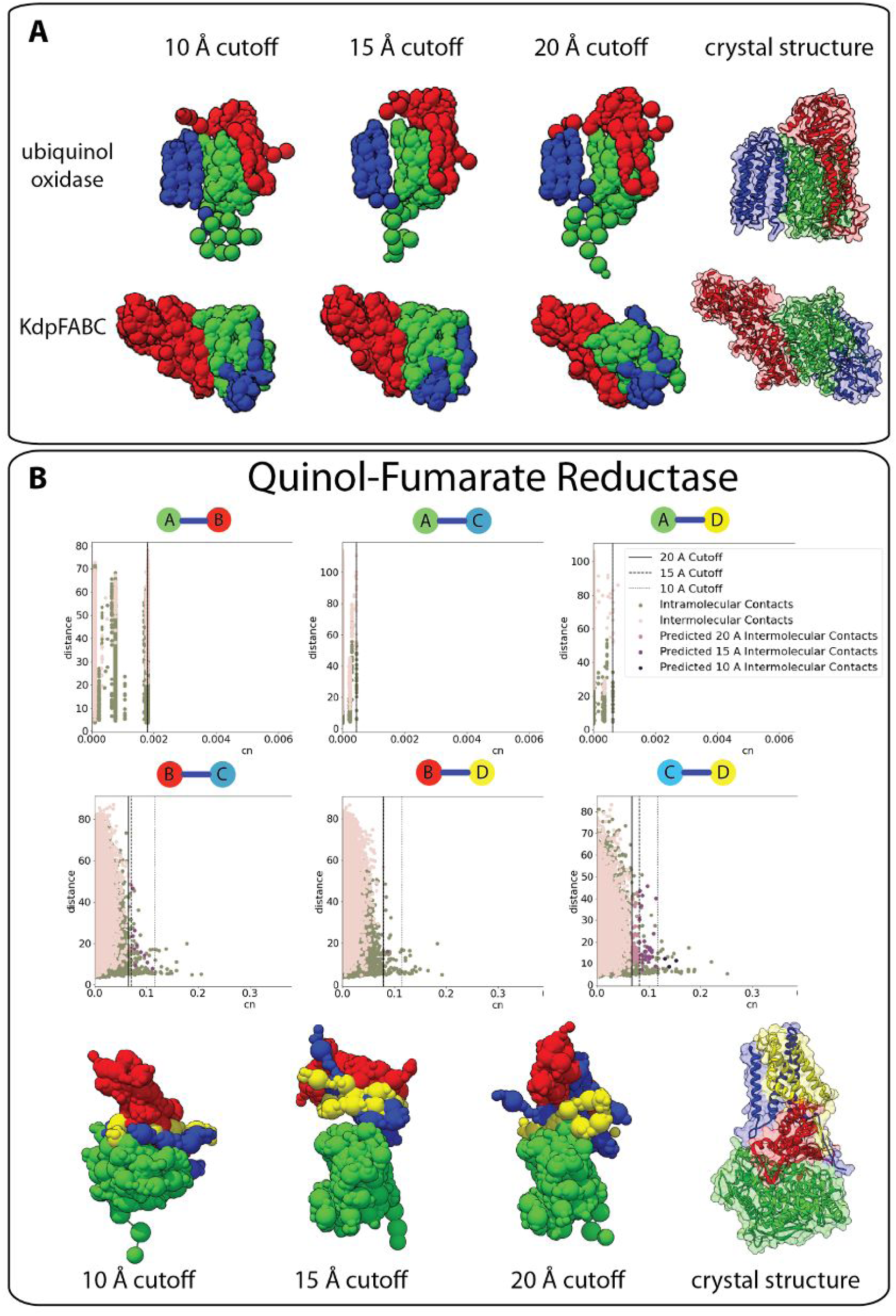
Evaluation of evolutionary coupling restraints on known structures. **A**. Different distance thresholds are tested for three complexes (PDBs: 1FFT, 5MRW, 1PRC) with varying degrees of evolutionary coupling coverage (details in supplemental materials). The IMP model ensembles are shown next to the crystal structures of the complexes. **B**. An example of a complex without sufficient evolutionary coupling coverage (PDB: 1L0V). The graphs show that subunit A has no couplings with any of the other subunits. The EM map combined with the other coupling information was not sufficient to produce an accurate ensemble of models.

We show the importance of including the evolutionary coupling distance restraint in our model for quinol-fumarate reductase (PDB: 1L0V) (**Figure 4B**). We were not able to detect any significant couplings between subunit A and any other subunit in the complex. This is further supported by the shape of the graphs, where there are no high scoring outliers that can be separated from the rest of the data (**Figure 4B**). In the final models, it is clear that the X-ray crystal structures, simulated EM map, and limited coupling did not provide enough spatial restraints to properly orient subunit A with respect to the remainder of the complex, and as a result, the remaining subunits struggled to find the global minimum that satisfied the restraints.

### Evolutionary couplings in modeling the dynamic BAM complex

To assess the ability of the evolutionary coupling restraints to capture protein complex dynamics, we built integrative models for the highly flexible *E. coli* BAM complex, again, using only X-ray crystal structures, evolutionary couplings, and 10-Å low-pass filtered PDB models to simulate EM maps. The BAM complex exists in a lateral-open state (PDB: 5D0Q) and an inward-open state (PDB: 5D0O) where the inward open state has conformational shifts of each protein and includes an additional subunit^39^. The conformational changes of each protein lead to slightly different intermolecular pairs. The conformational changes of subunits D and E lead to slightly different couplings being selected (**Figure 5**). Comparison of the structures show how these structural changes could result in different amino acid pairs due to the new intramolecular distances. Despite the structural shift and additional subunit, the evolutionary couplings integrated into the IMP scheme were able to predict the lateral-open state and inward-open state with sampling precisions of 21.7 Å and 9.0 Å, respectively.

**Figure 5:**
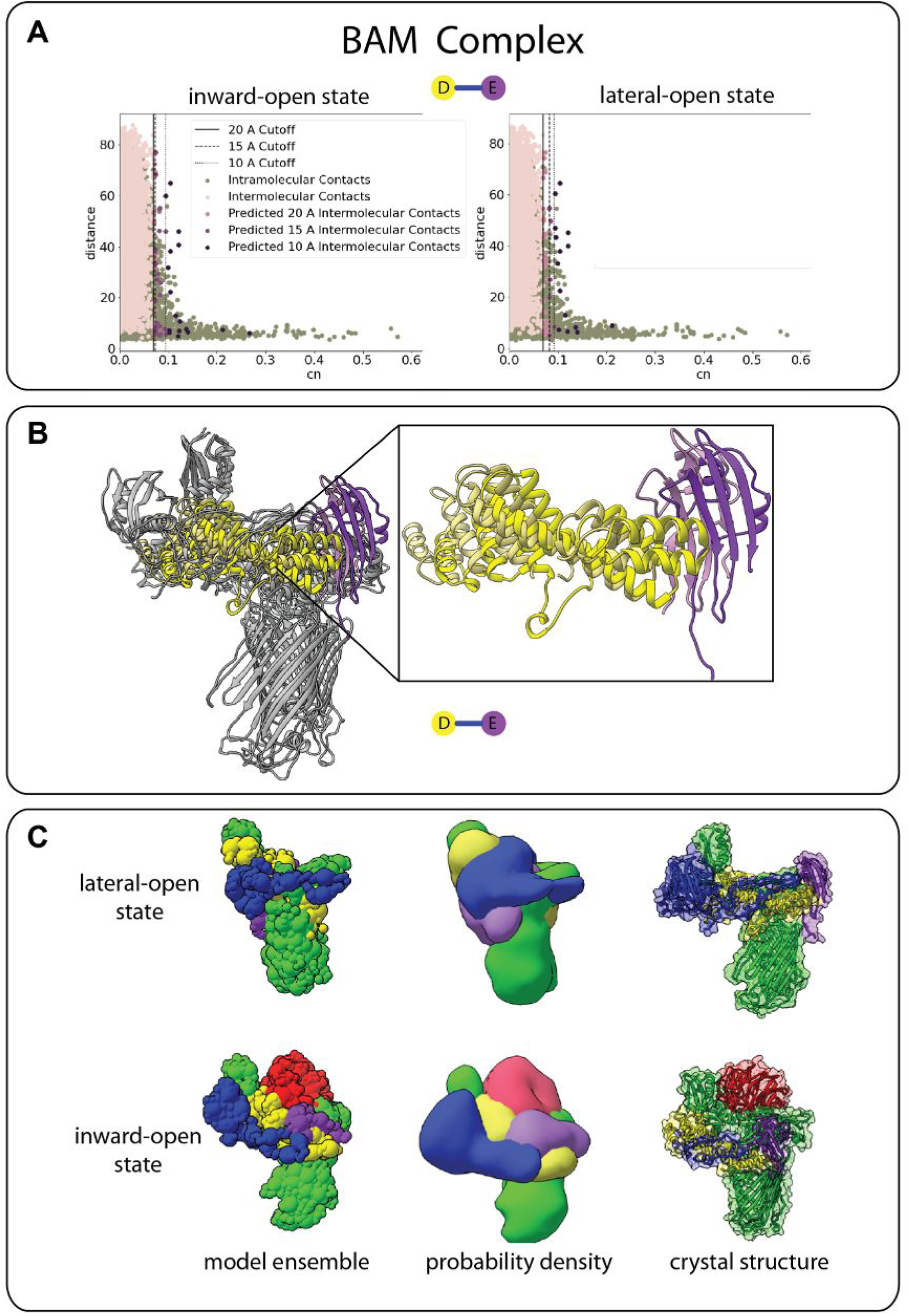
Evaluation of evolutionary coupling restraints on a dynamic structure. The BAM complex illustrates the robustness of evolutionary couplings against conformational changes in dynamic complexes. The BAM complex exists in a lateral-open state (PDB: 5D0Q) and an inward open state (PDB:5D0O), which includes an additional subunit. **A**. The internal control selected slightly different couplings based individual subunit conformational changes of BamD and BamE. **B**. The inward-open state is shown with the lighter yellow (BamD) and purple (BamE), outward the darker. **C**. The combination of the EM map and evolutionary couplings was enough data to produce each of the states as compared to the crystal structures.

### Integrative modeling of the bacterial holo-translocon using evolutionary couplings

After evaluating the accuracy of combining evolutionary distance restraints and EM maps, we were interested in using this method to systematically model the structure of the bacterial holo-translocon (HTL). The bacterial HTL is an assembly of three subcomplexes

--SecYEG, SecDF-YajC, and YidC. It is responsible for protein secretion across the membrane and membrane protein insertion. While partial structures are available for each of the subcomplexes, capturing the structure of the full complex is difficult due to the transient association of the subcomplexes^40^. While isolation of the complex from the membrane environment provides an additional challenge, advances in cryo-EM have made such tasks tangible.

Recently, a 14-Å resolution cryo-EM map for the bacterial HTL was published. While this map cannot be used for *de novo* model building, it still provides spatial information regarding the shape of the complex^35^. In addition, all subunits of the complex have X-ray crystal structures or could be modeled with high confidence, and we were able to use our internal calibration to extract evolutionary coupling between subunits to provide distance restraints between subunits. In our model, we treat each protein as a rigid body or chain of rigid bodies (for multi-domain proteins) and omit information regarding the subcomplexes that are formed prior to the HTL association (used later for validation). The central ensemble model and subunit probability densities were computed from 9,027 final models in the cluster (**Figure 6A**). The location of each of the subunits in our model agreed with that of a previous model built using the same EM map, SANS data, and biochemical information (**Supplementary Figure 1**), providing an independent confirmation of the model. The cross-correlation coefficient between the probability density and the cryo-EM map was 0.88.

**Figure 6:**
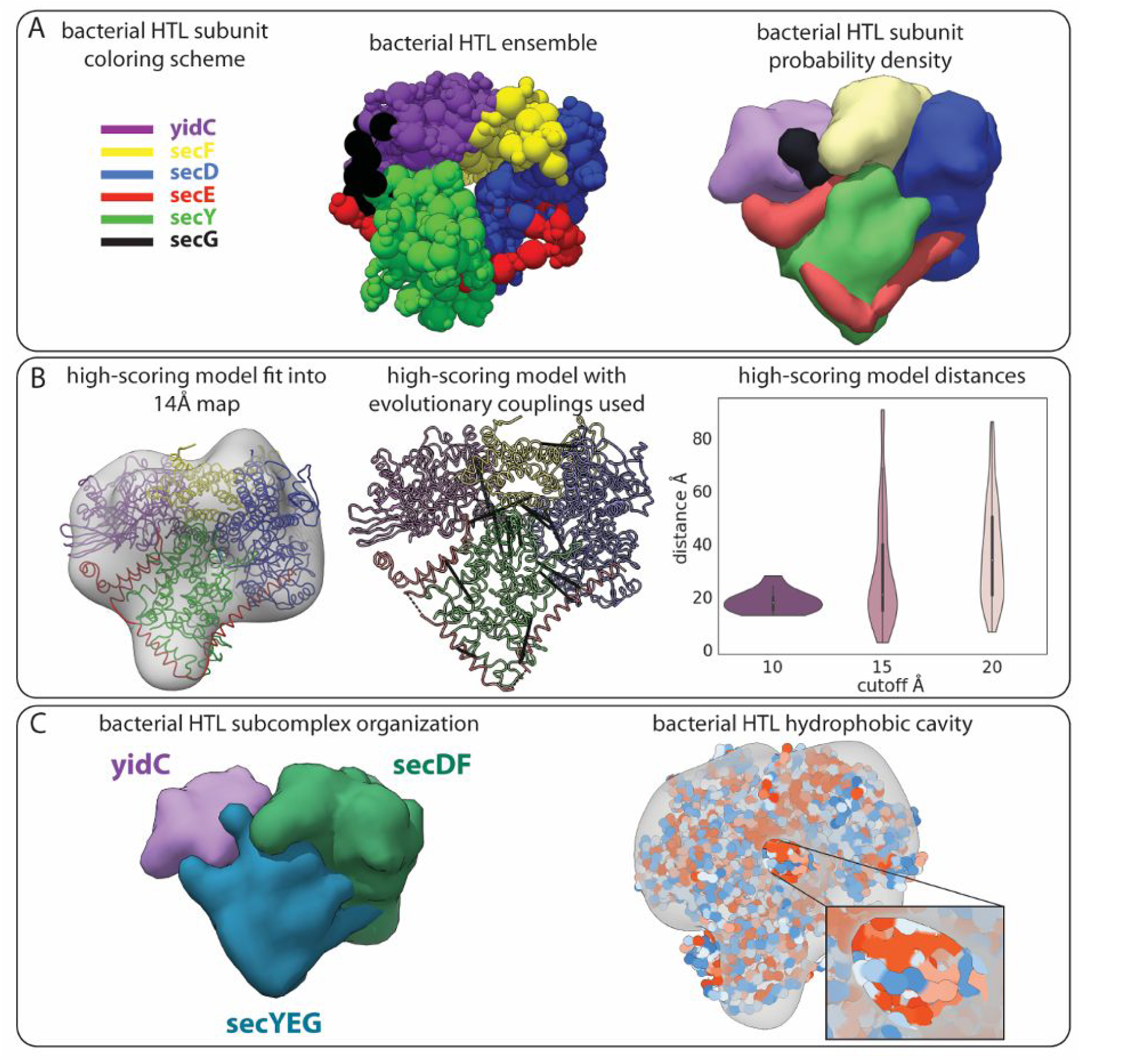
Integrative modeling of the bacterial holo-translocon. **A**. The model ensemble (middle) and subunit probability density (right) determined by integrative modeling using homology models, evolutionary coupling restraints, and an experimental 14-Å resolution cryo-EM map (EMDB: 3506). **B**. A high-scoring model from the ensemble is fit into the experimental map used for modeling (left). The evolutionary couplings used in modeling (10-Å distance threshold) plotted on the high-scoring model (middle). Intermolecular distances for each distance threshold (15- and 20-Å distances were not used in modeling) on the high-scoring model (right). **C**. The model agrees with other experimental information not used in sampling and analyzing the data. The probability density for the three subcomplexes are grouped together in our model (left). The hydrophobic surface is shown for the high-scoring model exposing the hydrophobic (red) cavity in the complex’s core (right).

In addition to validating our model against the cryo-EM map used, there is agreement between a high-scoring model from the final ensemble and the intermolecular evolutionary couplings used in the modeling (**Figure 6A**). We used the strictest threshold of 10 Å in selecting the score cutoffs for the model. However, many of the evolutionary couplings specific to the 15-Å and 20-Å cutoff groups were also satisfied in the model, even though they were not used in the construction of the model (**Figure 6B)**. We also see that at these lower score cutoffs there are a number of couplings that were not satisfied. This is consistent with the observation that the lower scores are closer to the center of the distribution, which is likely to have more false-positive couplings.

SecYEG is believed to form the membrane pore that works with YidC to insert hydrophobic protein segments into the lipid bilayer, which is accelerated by SecDF-yajC. When the probability densities for each of the subunits are combined into their prospective subcomplexes, we see a distinct localization of the subcomplexes, a feature that was not an input to the model (**Figure 6C**). A hydrophobic channel formed by the SecYEG and YidC subunits is evident in the high-scoring model (**Figure 6C**), where red indicates higher hydrophobicity and blue indicates higher hydrophilicity. This is consistent with the role of the sub-complexes in transporting hydrophobic polypeptides across the membrane^41^.

## Discussion

Here, we introduce an integration of evolutionary coupling data into an integrative modeling framework. Many previous reviews on integrative structural modeling methods have suggested the use of evolutionary coupling data would greatly benefit the models as a source of additional input information^42-44^. Our goal was to create a systematic method for selecting intermolecular couplings between proteins and utilize the residue couplings as a distance restraint in IMP. Because the evolutionary couplings are a source of co-evolutionary information, this method could greatly aid the fitting of crystal structures into medium- to low-resolution EM maps. We show this by modeling various protein complexes using only evolutionary coupling data paired with low-resolution EM maps. While we don’t explore the application of our evolutionary coupling distance restraint combined with other distances restraints (i.e. chemical crosslinks), we believe the couplings would only help to improve other experimental data sets.

In building the evolutionary coupling distance restraint, it was imperative that we had a universal method for selecting the residue pairs from the coupling data, and that we were somewhat judicious in doing so, as to allow for the appropriate selection and weighting of lower-scoring pairs. Previous use of evolutionary coupling data has shown to be effective in supplementing lower resolution data such as SAXS and EM maps, however, the selection criterion varies across studies^36; 37^. Our method allows for the recalibration of the score cutoff based on how well it fits the known, high confidence data. This internal calibration allows for us to select more pairs than a generalized score cutoff. In our test cases, we see a score cutoff as low as 0.027. We then balance this more tolerant selection method by weighting the evolutionary coupling used in modeling by their cn score, assuring that the highest-scoring pairs are prioritized over the lower. This increased tolerance selects a recommended score cutoff^17^ as compared to our three cutoffs --which vary for each pair (**Figure 1C**).

As a result of our selection method, we will likely have more false-positive couplings as compared to other more conservative score cutoffs. In the early days of residue co-evolution for protein-protein interactions, the goal was to maximize the precision of the prediction, which may be done by using a stricter cutoff. The authors, however, urge users to explore lower cutoffs^17^. Here, the use of lower cutoffs works well due to the scoring in IMP, where convergent high-scoring models are selected. In addition, in incorporating the evolutionary coupling distance restraint, there is no penalty for non-interacting proteins being in close spatial proximity. For the cases in which there is a false-positive pair with a high-score, the other couplings and the EM map help serve as a control.

While the co-evolutionary information stored in protein sequences is rich for predicting these pairwise contacts, it is important to note that the contact predictions are dependent on the depth of the sequence data. A limitation exists for those proteins that do not have sufficient coverage, as the recommended number of sequences is about one-third of the length of the sequence of interest. Nonetheless, there are now metagenomics databases^45^ with abundant information that may be used in combination with the sequences found on UniProt. The use of metagenomics in evolutionary couplings analyses has already been explored for building models of protein families with unknown structure and cannot be modeled using comparative modeling methods. This was able to generate models for 614 protein families with unknown structure^46^. While this has been used for intramolecular contacts and structure determination, we expect that the ability to predict couplings using metagenomics would carry over for the intermolecular couplings, such as with sequence data, and provide a supplemental source of sequence data for those proteins lacking sequence depth.

Using our method, we built an integrative model for the bacterial holo-translocon. While this model was built exclusively using the evolutionary coupling data at the 10-Å score cutoff and 14-Å resolution cryo-EM map, there are extensive experimental studies that support the model. The study that generated the cryo-EM map we used built a model for the complex using SANS data, the cryo-EM map, and biochemical information^35^. Our model agreed with this model on the localization of each subunit within the cryo-EM map (**Supplemental Figure 1**), however, because the full structure was not modeled, there were some areas of unassigned density in their map. Interestingly, in their study, they hypothesize that this unassigned density next to the SecYEG subcomplex corresponds to the N-terminal domains of SecE or YidC. In our model which has 100% sequence coverage, we indeed see the N-terminal of SecE occupying this area of unassigned density.

Our model is also supported by previous protein-protein interactions studies via chemical crosslinks and deletion analysis. Deletion studies in YidC have shown that YidC interacts directly with SecDF, in particular with SecF^47^. Our model not only places the SecDF subcomplex together, but it has YidC directly interacting with SecF. Additionally, these studies showed that residues 24-346 are required for YidC to interact with SecF, which is located on our interaction interface, and the mutant YidC with residues 527-548 deleted was still capable of binding SecF, a region that was not on the interaction interface of our model. Crosslink studies have also illuminated the interaction interface between YidC and SecY, another direct interaction we see in our model^48^. The strong cross-linked product residues of SecY had an average distance of 18 Å from YidC, while the weak cross-linked residues had an average distance of 17 Å from YidC and the non-cross-linked residues were 31 Å from YidC (all between Ca atoms). Taking these various independent experiments together, they provide us with confidence in our model.

In this work, we have developed a distance restraint based on evolutionary coupling data to be used in IMP. We have rigorously benchmarked appropriate thresholds that account for noise in the data through an internal score calibration. We have tested this approach on complexes of varying complexity including complexes with conformational changes and additional subunits. Finally, we applied this method to building an integrative model of the bacterial holo-translocon that agrees with previous experimental data. We believe that because this restraint is built using co-evolutionary information and does not require any additional experiments, it will be useful for assembling atomic structures into medium- to low-resolution EM maps, providing additional restraints to integrative models when required.

## Acknowledgments

All code and sample files are available at https://github.com/marcottelab/CoEVxIMP, along with our final integrative model of the bacterial holo-translocon. We are deeply grateful to A. Šali, R. Ramachandran, and B. Webb for assistance with IMP and helpful feedback and comments on this manuscript. This work was supported in part by Welch Foundation Research Grants F-1938 (to D.W.T.) and F-1515 (to E.M.M.), Army Research Office Grant W911NF-15-1-0120 (to D.W.T.), a Robert J. Kleberg, Jr., and Helen C. Kleberg Foundation Medical Research Award (to D.W.T.), and grants from the National Institutes of Health GM122480, DK110520, and HD085901 (to E.M.M.) and R35GM138348 (to D.W.T.). C.L.M is an NSF Graduate Research Fellow supported by the National Science Foundation (2019238253). D.W.T is a CPRIT Scholar supported by the Cancer Prevention and Research Institute of Texas (RR160088) and an Army Young Investigator supported by the Army Research Office (W911NF-19-1-0021).

## Figure legends

**Supplemental Figure 1.**
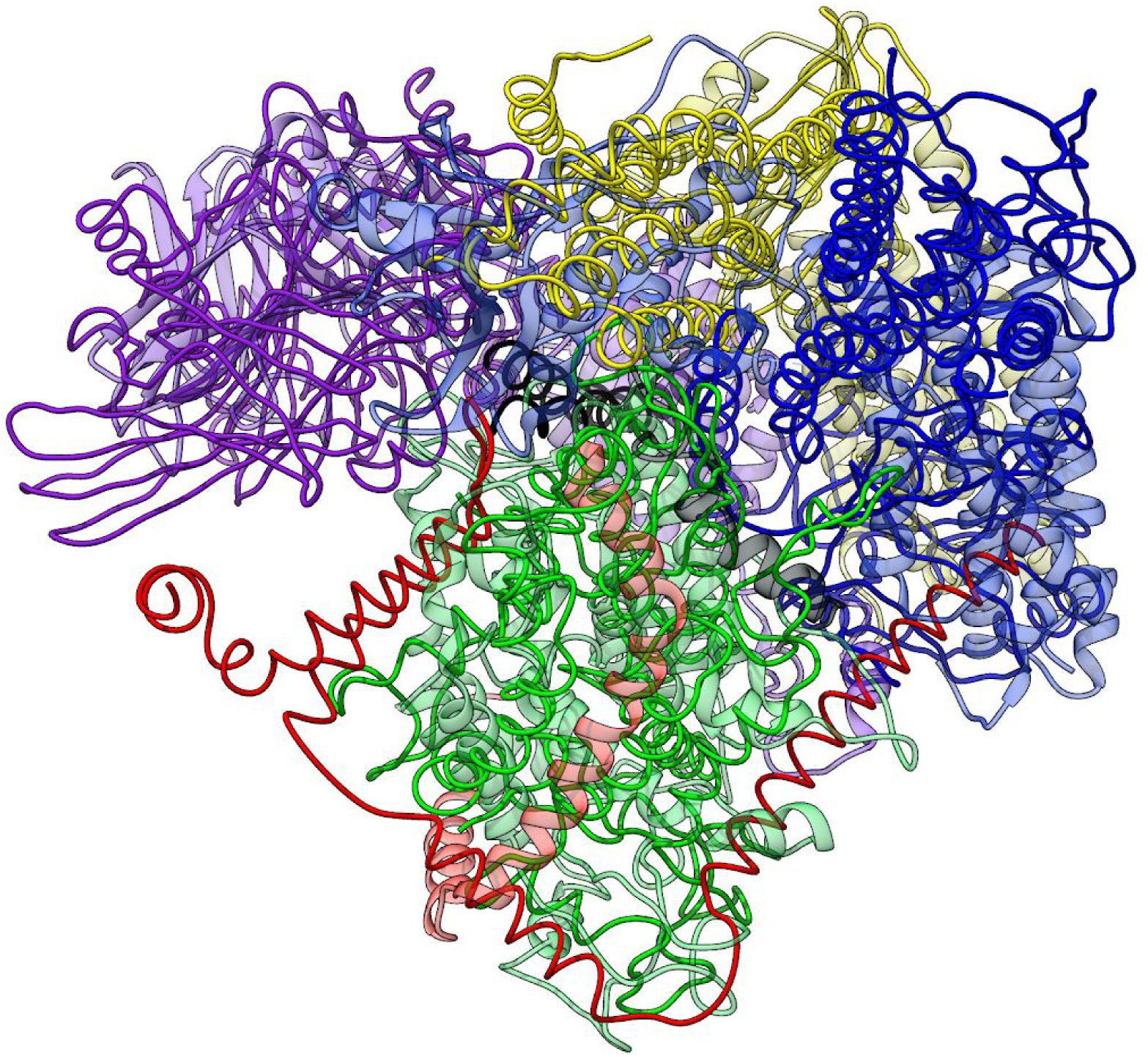
Integrative model of the bacterial holo-translocon superimposed on the published model (PDB: 5MG3).

**Supplemental Table 1.**
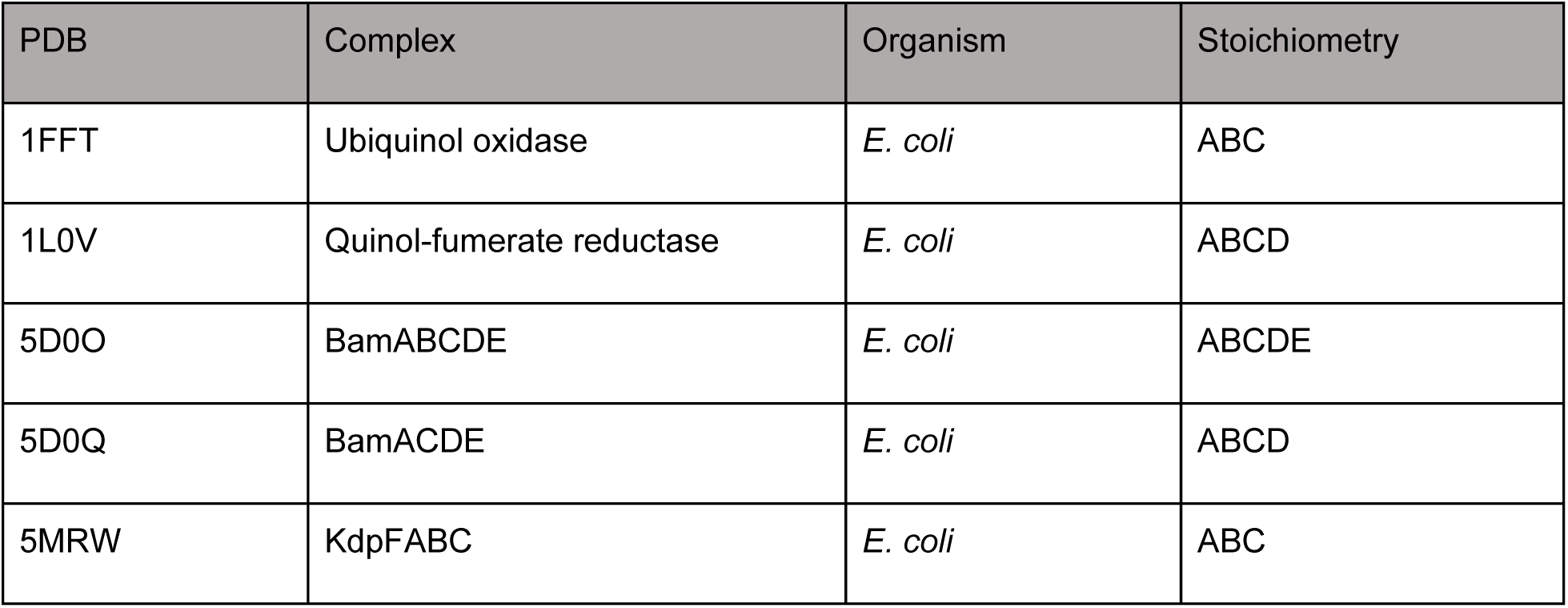
Complexes modeled.

## References

1. Uniprot: The universal protein knowledgebase. 2017. Nucleic Acids Res. 45(D1):D158–D169.

2. Katoh K, Misawa K, Kuma Ki, Miyata T. 2002. Mafft: A novel method for rapid multiple sequence alignment based on fast fourier transform. Nucleic Acids Res. 30(14):3059–3066.

3. Notredame C, Higgins DG, Heringa J. 2000. T-coffee: A novel method for fast and accurate multiple sequence alignment. J Mol Biol. 302(1):205–217.

4. Do CB, Mahabhashyam MS, Brudno M, Batzoglou S. 2005. Probcons: Probabilistic consistency-based multiple sequence alignment. Genome Res. 15(2):330–340.

5. Brown DP, Krishnamurthy N, Sjölander K. 2007. Automated protein subfamily identification and classification. PLoS Comput Biol. 3(8):e160.

6. Brandt BW, Feenstra KA, Heringa J. 2010. Multi-harmony: Detecting functional specificity from sequence alignment. Nucleic Acids Res. 38(Suppl_2):W35–W40.

7. Rausell A, Juan D, Pazos F, Valencia A. 2010. Protein interactions and ligand binding: From protein subfamilies to functional specificity. Proceedings of the National Academy of Sciences. 107(5):1995–2000.

8. Morcos F, Pagnani A, Lunt B, Bertolino A, Marks DS, Sander C, Zecchina R, Onuchic JN, Hwa T, Weigt M. 2011. Direct-coupling analysis of residue coevolution captures native contacts across many protein families. Proceedings of the National Academy of Sciences. 108(49):E1293–E1301.

9. Göbel U, Sander C, Schneider R, Valencia A. 1994. Correlated mutations and residue contacts in proteins. Proteins: Structure, Function, and Bioinformatics. 18(4):309–317.

10. Marks DS, Colwell LJ, Sheridan R, Hopf TA, Pagnani A, Zecchina R, Sander C. 2011. Protein 3d structure computed from evolutionary sequence variation. PLoS One. 6(12):e28766.

11. Marks DS, Hopf TA, Sander C. 2012. Protein structure prediction from sequence variation. Nat Biotechnol. 30(11):1072.

12. Senior AW, Evans R, Jumper J, Kirkpatrick J, Sifre L, Green T, Qin C, Žídek A, Nelson AW, Bridgland A. 2020. Improved protein structure prediction using potentials from deep learning. Nature. 577(7792):706–710.

13. Huang YJ, Brock KP, Sander C, Marks DS, Montelione GT. 2018. A hybrid approach for protein structure determination combining sparse nmr with evolutionary coupling sequence data. Integrative structural biology with hybrid methods. Springer. p. 153–169.

14. Huang YJ, Brock KP, Ishida Y, Swapna GV, Inouye M, Marks DS, Sander C, Montelione GT. 2019. Combining evolutionary covariance and nmr data for protein structure determination. Methods enzymol. Elsevier. p. 363–392.

15. Hopf TA, Ingraham JB, Poelwijk FJ, Schärfe CP, Springer M, Sander C, Marks DS. 2017. Mutation effects predicted from sequence co-variation. Nat Biotechnol. 35(2):128–135.

16. Weinreb C, Riesselman AJ, Ingraham JB, Gross T, Sander C, Marks DS. 2016. 3d rna and functional interactions from evolutionary couplings. Cell. 165(4):963–975.

17. Hopf TA, Schärfe CP, Rodrigues JP, Green AG, Kohlbacher O, Sander C, Bonvin AM, Marks DS. 2014. Sequence co-evolution gives 3d contacts and structures of protein complexes. Elife. 3:e03430.

18. Ovchinnikov S, Kamisetty H, Baker D. 2014. Robust and accurate prediction of residue–residue interactions across protein interfaces using evolutionary information. Elife. 3:e02030.

19. Cong Q, Anishchenko I, Ovchinnikov S, Baker D. 2019. Protein interaction networks revealed by proteome coevolution. Science. 365(6449):185–189.

20. Green AG, Elhabashy H, Brock KP, Maddamsetti R, Kohlbacher O, Marks DS. 2019. Proteome-scale discovery of protein interactions with residue-level resolution using sequence coevolution. bioRxiv.791293.

21. Webb B, Viswanath S, Bonomi M, Pellarin R, Greenberg CH, Saltzberg D, Sali A. 2018. Integrative structure modeling with the integrative modeling platform. Protein Sci. 27(1):245–258.

22. Shi Y, Fernandez-Martinez J, Tjioe E, Pellarin R, Kim SJ, Williams R, Schneidman-Duhovny D, Sali A, Rout MP, Chait BT. 2014. Structural characterization by cross-linking reveals the detailed architecture of a coatomer-related heptameric module from the nuclear pore complex. Mol Cell Proteomics. 13(11):2927–2943.

23. Ganesan SJ, Feyder MJ, Chemmama IE, Fang F, Rout MP, Chait BT, Shi Y, Munson M, Sali A. 2020. Integrative structure and function of the yeast exocyst complex. Protein Sci.

24. Pintilie G, Chiu W. 2012. Comparison of segger and other methods for segmentation and rigid-body docking of molecular components in cryo-em density maps. Biopolymers. 97(9):742–760.

25. Lander GC, Estrin E, Matyskiela ME, Bashore C, Nogales E, Martin A. 2012. Complete subunit architecture of the proteasome regulatory particle. Nature. 482(7384):186–191.

26. Hopf TA, Green AG, Schubert B, Mersmann S, Schärfe CP, Ingraham JB, Toth-Petroczy A, Brock K, Riesselman AJ, Palmedo P. 2019. The evcouplings python framework for coevolutionary sequence analysis. Bioinformatics. 35(9):1582–1584.

27. Suzek BE, Huang H, McGarvey P, Mazumder R, Wu CH. 2007. Uniref: Comprehensive and non-redundant uniprot reference clusters. Bioinformatics. 23(10):1282–1288.

28. Hockenberry AJ, Wilke CO. 2019. Evolutionary couplings detect side-chain interactions. PeerJ. 7:e7280.

29. Saltzberg D, Greenberg CH, Viswanath S, Chemmama I, Webb B, Pellarin R, Echeverria I, Sali A. 2019. Modeling biological complexes using integrative modeling platform. Biomolecular simulations. Springer. p. 353–377.

30. Kawabata T. 2018. Gaussian-input gaussian mixture model for representing density maps and atomic models. J Struct Biol. 203(1):1–16.

31. Viswanath S, Chemmama IE, Cimermancic P, Sali A. 2017. Assessing exhaustiveness of stochastic sampling for integrative modeling of macromolecular structures. Biophys J. 113(11):2344–2353.

32. McInnes L, Healy J, Astels S. 2017. Hdbscan: Hierarchical density based clustering. Journal of Open Source Software. 2(11):205.

33. Roy A, Kucukural A, Zhang Y. 2010. I-tasser: A unified platform for automated protein structure and function prediction. Nat Protoc. 5(4):725.

34. Tanaka Y, Izumioka A, Hamid AA, Fujii A, Haruyama T, Furukawa A, Tsukazaki T. 2018. 2.8-å crystal structure of escherichia coli yidc revealing all core regions, including flexible c2 loop. Biochemical and biophysical research communications. 505(1):141–145.

35. Botte M, Zaccai NR, Martin R, Knoops K, Papai G, Zou J, Deniaud A, Karuppasamy M, Jiang Q, Roy AS. 2016. A central cavity within the holo-translocon suggests a mechanism for membrane protein insertion. Sci Rep. 6(1):1–13.

36. Leone V, Faraldo-Gómez JD. 2016. Structure and mechanism of the atp synthase membrane motor inferred from quantitative integrative modeling. J Gen Physiol. 148(6):441–457.

37. Zhao C, Shukla D. 2018. Saxs-guided enhanced unbiased sampling for structure determination of proteins and complexes. Sci Rep. 8(1):1–13.

38. Bürmann F, Shin H-C, Basquin J, Soh Y-M, Giménez-Oya V, Kim Y-G, Oh B-H, Gruber S. 2013. An asymmetric smc–kleisin bridge in prokaryotic condensin. Nat Struct Mol Biol. 20(3):371.

39. Gu Y, Li H, Dong H, Zeng Y, Zhang Z, Paterson NG, Stansfeld PJ, Wang Z, Zhang Y, Wang W. 2016. Structural basis of outer membrane protein insertion by the bam complex. Nature. 531(7592):64–69.

40. Duong F. 2014. Capturing the bacterial holo-complex. Proceedings of the National Academy of Sciences. 111(13):4739–4740.

41. Sachelaru I, Winter L, Knyazev DG, Zimmermann M, Vogt A, Kuttner R, Ollinger N, Siligan C, Pohl P, Koch H-G. 2017. Yidc and secyeg form a heterotetrameric protein translocation channel. Sci Rep. 7(1):1–15.

42. Braitbard M, Schneidman-Duhovny D, Kalisman N. 2019. Integrative structure modeling: Overview and assessment. Annual review of biochemistry. 88.

43. van Zundert GC, Melquiond AS, Bonvin AM. 2015. Integrative modeling of biomolecular complexes: Haddocking with cryo-electron microscopy data. Structure. 23(5):949–960.

44. Joseph AP, Polles G, Alber F, Topf M. 2017. Integrative modelling of cellular assemblies. Curr Opin Struct Biol. 46:102–109.

45. Mitchell AL, Almeida A, Beracochea M, Boland M, Burgin J, Cochrane G, Crusoe MR, Kale V, Potter SC, Richardson LJ. 2020. Mgnify: The microbiome analysis resource in 2020. Nucleic Acids Res. 48(D1):D570–D578.

46. Ovchinnikov S, Park H, Varghese N, Huang P-S, Pavlopoulos GA, Kim DE, Kamisetty H, Kyrpides NC, Baker D. 2017. Protein structure determination using metagenome sequence data. Science. 355(6322):294–298.

47. Xie K, Kiefer D, Nagler G, Dalbey RE, Kuhn A. 2006. Different regions of the nonconserved large periplasmic domain of escherichia coli yidc are involved in the secf interaction and membrane insertase activity. Biochemistry. 45(44):13401–13408.

48. Sachelaru I, Petriman NA, Kudva R, Kuhn P, Welte T, Knapp B, Drepper F, Warscheid B, Koch H-G. 2013. Yidc occupies the lateral gate of the secyeg translocon and is sequentially displaced by a nascent membrane protein. J Biol Chem. 288(23):16295–16307.

